# Tissue specific *de novo* transcriptome sequencing and expression analysis of candidate genes for terpenoid biosynthesis in *Piper longum*

**DOI:** 10.1101/2025.10.29.685027

**Authors:** Manaswini Dash, Bhaskar Chandra Sahoo, Suprava Sahoo, Basudeba Kar

## Abstract

*Piper longum* L. an economically important plant lacks adequate molecular resources towards unmasking molecular mechanisms for terpenoid biosynthesis. The transcriptome profiling generated 36.91 and 36.24 million raw reads in leaf and fruit tissue respectively. The *de novo* transcriptome assembly was performed with 36,688,460 and 36,011,305 clean reads, which subsequently produced 40,033 and 44,025 coding sequences (CDS) in leaf and fruit samples, respectively. The CDS were annotated using different databases such as Gene Ontology (GO) and Kyoto Encyclopedia of Genes and Genomes (KEGG). These annotations resulted mining of various trait specific genes including important terpenoid biosynthesis genes. Presence of 53 and 58 terpenoid compounds in essential oils of leaf and fruit of long pepper have also been identified using GC-MS. Additionally, relative gene expression of eight terpenoid biosynthesis genes each for both leaf and fruit were conducted using real-time polymerase chain reaction (RT-PCR). For further genetic diversity study, 4085 and 4564 number of SSRs were identified in leaf and fruit, respectively. This study significantly contributes to the understanding of long pepper at the molecular level by expanding the available genetic resources. This paves the way to improve long pepper varieties and develop new genotype with desirable quality traits.

## Introduction

*Piper longum* L. (Indian long pepper) belongs to the family Piperaceae and is one of the best-known species for its enormous medicinal properties and economic importance. It is an aromatic, creeping perennial herb widely distributed in tropical and subtropical regions around the globe, throughout the Indian subcontinent, Sri Lanka, and Middle Eastern countries (Sarwar et al., 2015). *P. longum* is generally cultivated for its fruits, commonly known as Pippali (Scott et al., 2008). The dried fruits are generally used as an Indian dietary spice as well as for treating stomach disorders and analgesia in Traditional Chinese Medicine (Abdubakiev et al., 2019). In addition, long pepper fruits have been utilized in the Ayurvedic medicinal system to treat several respiratory tract diseases like cough, asthma, cold, and bronchitis. Furthermore, the fruits are also used for the alleviation of gastrointestinal disorders, metabolic imbalance, spleen disorders, arthritic complications, tuberculosis, and jaundice (Liu et al., 2011). Besides the fruit, leaf is also traditionally used in treating several ailments (Sultana et al., 2019). The leaves of this species possess bioactivities such as antioxidant, antibacterial, antifungal, larvicidal, ovicidal, antidermaptophytic, and anthelmintic (Dash et al., 2022, Dey et al., 2020, Das et al., 2012, D’Cruz et al., 1980). Due to its wide range of ethnomedicinal, ayurvedic, and pharmaceutical applications, Pippali is in high demand in domestic as well as global markets.

The long pepper fruit and leaf volatiles are mainly comprised of monoterpenes like α-pinene, *α*-thujene, camphene, *β*-pinene, limonene, linalool, camphor, myrcene, (*E*)-*β*-ocimene, 2- heptyl acetate, trans-*β*-terpineol, *γ*-terpinene, terpinolene, *cis*-carveol and sesquiterpenes like germacrene-D, *E*-nerolidol, *Z*-caryophyllene, 8-heptadecene, *α*-muurolol, *α*-humulene, n-pentadecane, *γ*-cadinene, *n*-heptadecane, *n*-hexadecanol, *n*-nonadecane, 9-nonadecene along with certain diterpenes such as undecanoic acid, phytol, untriacontane, heptacosane and n-eicosane (Dash et al., 2022, Hieu et al. 2018, Varughese et al., 2016, Rameshkumar et al., 2011, Bhuiyan et al., 2013, Liu et al., 2007, Tewtrakul et al., 2000). The two universal precursor molecules; isopentyl pyrophosphate (IPP) for cytosolic mevalonic acid (MVA) pathway and dimethylallyl pyrophosphate (DMAPP) for plastidial methylerythritol phosphate/ 1-deoxy-D-xylulose 5-phosphate (MEP/DXP) pathway are used by plants to synthesize the backbone of terpenoids. IPP and DMAPP undergo several enzymatic reactions to form linear prenyl diphosphates such as; geranyl diphosphate (GPP), farnesyl diphosphate (FPP), geranylgeranyl diphosphate (GGPP), and farnesyl geranyl diphosphate (FGPP). The precursors GPP, FPP, GGPP, and FGPP are involved in the biosynthesis of different classes of terpenoids by various terpene synthase (TPS) enzymes (Abdallah et al., 2017).

Nevertheless, no reports are available on the genes involved in the terpenoid biosynthesis of long pepper plant. Despite having enormous medicinal and economic values, the availability of genomic information is minimal. The lack of genetic resources circumscribes the identification and characterization of novel genes regulating different pathways. There are few ESTs (Expressed Sequenced Tags) sequences available in long pepper in NCBI (National Centre for Biotechnology Information) databases, due to which it will be tough to map the geographical diversification of *P. longum*.

Considering these constraints, trait-specific genes implicated in a wide range of cellular, molecular, and physiological processes were mined by a *de novo* transcriptome study of long pepper. In addition, the current study aimed to generate high quality EST-SSRs (Expressed Sequence Tag-derived Simple Sequence Repeats) from the clean transcripts. Putative genes involved in terpenoid biosynthesis in long pepper were identified by analysing the transcriptome profile. An expression study of five terpenoid synthesis genes was carried out using Real Time PCR (RT-PCR) for further validation. These data will provide valuable resources for future investigations into long pepper’s genetic diversity, structural, and functional genomics.

## Materials and methods

### Sample collection

*Piper longum* plant was collected from Patbil, Mayurbhanj, north central plateau agroclimatic zone of Odisha (Latitude: 21° 38ʹ 55ʺ N; Longitude: 86° 14ʹ 56ʺ E; Altitude: 928 m) in the month of February 2020. The plant was identified and authenticated by renowned taxonomist and deposited in the herbarium of Centre for Biotechnology, Siksha O Anusandhan Deemed to be University, Odisha, India (20° 17ʹ 4.5852ʺ N, 85° 46ʹ 30.8496ʺ E) bearing voucher specimen number 2254/CBT. Both leaf and fruit samples of the plant were taken for RNA sequencing.

### RNA isolation, library construction and Illumina sequencing

The young leaves and fruits were collected in triplicates. Both the plant materials were washed properly with distilled water and then with 0.1% Diethyl pyrocarbonate (DEPC) water. The samples were then chopped and immediately snap frozen with liquid nitrogen and followed by storing at -80° C until further use. Total RNA was isolated using RNeasy mini kit and subjected to on column DNase treatment (Qiagen, Hilden, Germany) following manufacturer’s instructions. The quality and quantity of total RNA was estimated using Nanodrop spectrophotometer (Thermo Fisher Scientific; 2000) and Qubit RNA HS assay kit (Q32855). The RNA samples with 260/280 and 260/230 nm ratio ranging from 2.0-2.2 and RNA integrity number (RIN) more than 8.0 were assessed on Agilent 2200 TapeStation for construction of cDNA library.

The mRNA libraries were prepared using Illumina-compatible NEBNext^®^ Ultra^TM^ II Directional RNA Library Prep Kit (New England Biolabs, MA, USA). The mRNA molecules containing poly A were isolated using magnetic beads linked to poly T oligonucleotides. Poly A mRNAs were first fragmented followed by primed with random hexamer primers prior to cDNA (complementary DNA) synthesis. The first strand cDNA was generated followed by the second strand cDNA synthesis. The double stranded cDNAs were end-repaired and adenylated at 3’ ends to avoid low rate of chimera formation. Purification of the double stranded cDNAs were done using NEBNext sample purification beads. The 3’ adenylation end cDNAs were then ligated to Illumina adapters followed by 15 cycles of PCR (Polymerase Chain Reaction) amplification to enrich the library products. The obtained PCR products were purified followed by library quality control check. A Qubit fluorometer (Thermo Fisher Scientific, MA, USA) was used to quantify Illumina-compatible sequencing libraries, and an Agilent 2200 TapeStation was used to analyze the fragment size distribution. The band at around 447 bp confirmed the high level of purity. The libraries were then subjected to 150 paired end sequencing on Illumina HiSeq×Ten sequencer (Illumina, San Diego, USA) according to manufacturer’s instructions.

### *De novo* assembly and CDS identification

The raw sequence data obtained during sequencing were assessed for quality check, which included trimming of adaptor or primer sequences using FastQC software (version 0.11.8) (Andrews, 2010). Furthermore, the reads were pre-processed filtering off the low-quality reads having base Q value < 30 (TrimGalore tool) (Raghavan et al., 2022). The generated clean reads were assembled employing a graph-based approach by RnaSPAdes software (version 3.9.1) (Bankevich et al., 2012) with a k-mer size of 55. The non-redundant (NR) clustered transcripts (unigenes) were predicted by using CD-HIT software (version 4.6) (Li et al., 2006) by eliminating shorter redundant sequences with 95% similarity threshold. Finally, the TransDecoder tool (https://github.com/TransDecoder/TransDecoder/wiki) was used to filter out only the coding sequences (CDS) from all the assembled unigenes of leaf and fruit samples.

### Identification of homologous sequences and functional annotation of unigenes

Annotating the transcripts with information from a variety of databases based on a homology-based technique could aid in accurate annotation and, in turn, provide insight into the functioning of anything. Annotation in this report was obtained from several sources, including functional annotation against Viridiplantae protein sequences from Uniprot Protein Database (e value≤ 10^-5^) and Gene Ontology (GO) mapping of NR annotated sequences using Diamond BLAST. Further, transcripts were compared to the Kyoto Encyclopedia of Genes and Genomes (KEGG) using the KEGG automatic annotation system (KAAS) (Moriya et al., 2007) to identify orthologs and predict metabolic pathways in *P. longum*.

### Essential oil extraction and GC-MS analysis

The dried leaves (100g) and fruits (100g) of *P. longum* were taken, crushed, and subjected for essential oil (EO) extraction by hydro-distillation method using a Clevenger apparatus. After a period of 4 hours, the oil yield of leaf and fruit was determined based on the dry weight (v/w). The obtained EOs were then treated with anhydrous sodium sulfate (Na_2_SO_4_) for the removal of moisture traces and stored in glass vials at 4°C until use. The terpenoid constituents of *P. longum* leaves and fruits were analyzed by gas chromatography and mass spectrometry (GC-MS) using Clarus 580 Gas Chromatograph (Perkin Elmer, USA) attached to SQ8S Mass Detector. Elite-5 MS capillary column of length 30 m× 0.25 mm internal diameter and film thickness of 0.25 µm was installed in the instrument. The mobile phase carried Helium gas at a flow rate of 1.0 mL/min. The neat EOs of 0.1 µl of both leaf and fruit were injected individually with a temperature profile set initially at 60°C followed by ramping at 4°C/min to 120°C, then at 2°C/min to 180°C and finally at 5°C/min up to 250°C with 1 min hold. The inlet line temperature and source temperature were set at 250 °C and 150 °C respectively. Mass scanning of 50-600 m/z and 70eV electron ionization (EI) energy were applied. The ion chromatogram and mass spectra were obtained from the Turbo Mass software (Perkin Elmer Inc., USA; Version 6.1.0). For determining retention index (RI) of the constituents, homologous n-alkane series (C_8_-C_20_) (Sigma Aldrich, USA) with same operating conditions. Identification of compounds were done by comparing the mass spectra with the inbuilt NIST/EPA/NIH mass spectra library (NIST, 2020) followed by further validation of the RI values acquired from the experiment with those published in the literature and Adams library (Adams, 2007).

### Mining of SSRs

In order to extract SSRs (Simple Sequence Repeats) from RNA-sequence transcripts of both leaf and fruit tissues, Perl script of MIcro SAtellite (MISA) identification programme (http://pgrc.ipk-gatersleben.de/misa/) has been used. Mono-, di-, tri-, tetra-, penta-, and hexanucleotide repeats, as well as compound microsatellites, were analysed. The minimum number of repetitions for mono-, di-, tri-, tetra-, penta-, and hexanucleotide were set to 10, 6, 5, 3, 5, and 5 respectively for both leaf and fruit samples.

### RNA extraction and cDNA synthesis

Total RNAs were isolated from leaf and fruit of the long pepper plant using TRIzol (Invitrogen) according to the manufacturer’s instructions. Afterwards, genomic DNA contamination was removed via purification with DNaseI, RNase-free (Thermo Fisher Scientific, MA, USA) from the extracted total RNAs. The quantities of RNAs were determined by Nanodrop spectrometer (BioSpectrometer, Eppendorf AG, Hamburg, Germany). Purified RNAs (500 ng) were reverse transcribed using GoScript^TM^ Reverse transcription system (Promega, USA) to synthesize cDNA for both leaf and fruit.

### Real time PCR relative gene expression analysis

In order to experimentally validate the putative transcripts of long pepper that are involved in terpene biosynthesis, RT-PCR was carried out. The gene expression was investigated by designing eight terpene biosynthesis genes from the immaculate transcripts each for both leaf and fruit tissues. Total eighteen number of genes; germacrene-D synthase (1), caryophyllene synthase (2), humulene synthase (2), copaene synthase (1), *δ*-cadinene synthase (2), cadinol synthase (2), *α*-terpineol synthase (2), terpene synthase (2), monoterpene synthase (2), sesquiterpene synthase (2) were chosen for studying the relative gene expression (Table 3.3). As endogenous reference, actin (2) and ubiquitin (2) genes were used. The experiment was executed in a 10 µl reaction system consisted of PowerUp^TM^ SYBR^TM^ Green Master mix (5 µl), cDNA (1 µl), forward and reverse primers (5 pM each) (1 µl), and nuclease-free water (2 µl) using Applied Biosystems StepOnePlus^TM^ RT-PCR system. The reaction was amplified by cycling the temperature as follows: 95° C for 30 seconds, 40 cycles of 95° C for 15 seconds and 60° C for 1 minute. To further confirm the reaction’s specificity, a melt curve was run after the assay. Three independent biological replicates and three technical replicates of each RT- PCR experiment was performed and the relative gene expression was determined by comparative CT (cycle threshold) method (Livak and Schmittgen, 2001).

## Results

### Sequencing and *de novo* assembly

To know the scientific mechanism of terpene biosynthesis in long pepper plant, the transcriptomic sequencing was carried out. After transcriptomic sequencing, the sequences were processed for quality check. A total of 36,914,662 (36.91 million) and 36,244,013 (36.24 million) raw reads were generated in leaf and fruit samples, respectively from Illumina HiSeq×10 sequencer, accounting for approximately 4.6 Gb.

The FastQC software was used to perform quality control analysis on the raw reads. After eliminating the adaptor and filtering off the low-quality sequences from the raw reads, a total of 36,688,460 (36.69 million) (99.39%) and 36,011,305 (36.01 million) (99.36%) high quality reads have been generated for long pepper leaf and fruit respectively (Table 2, Fig. 1). Further, only high-quality reads without ambiguous “N” were confirmed.

**Fig 1.**
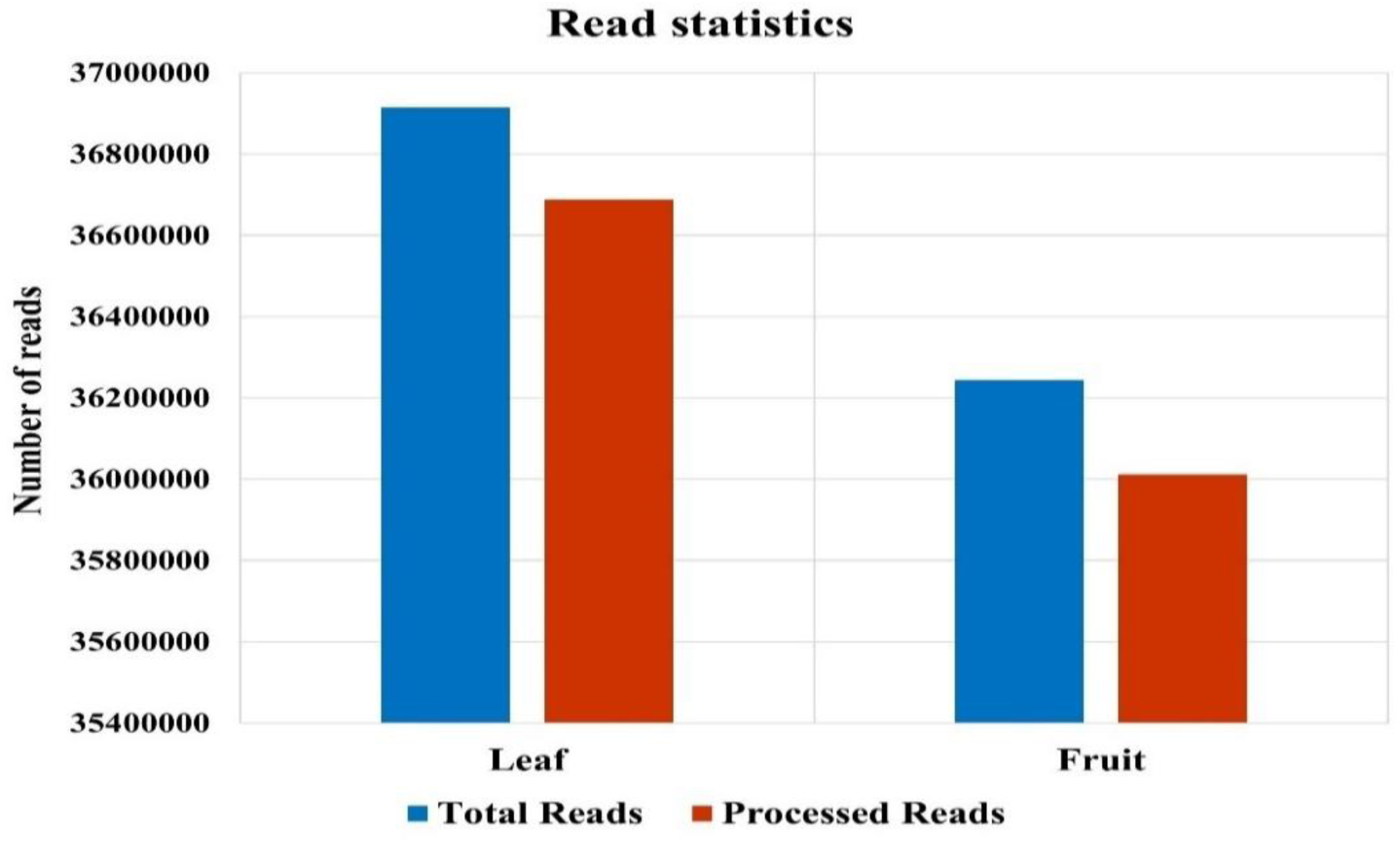
Read statistics of leaf and fruit samples of *P*. *longum*.

**Table 1:**
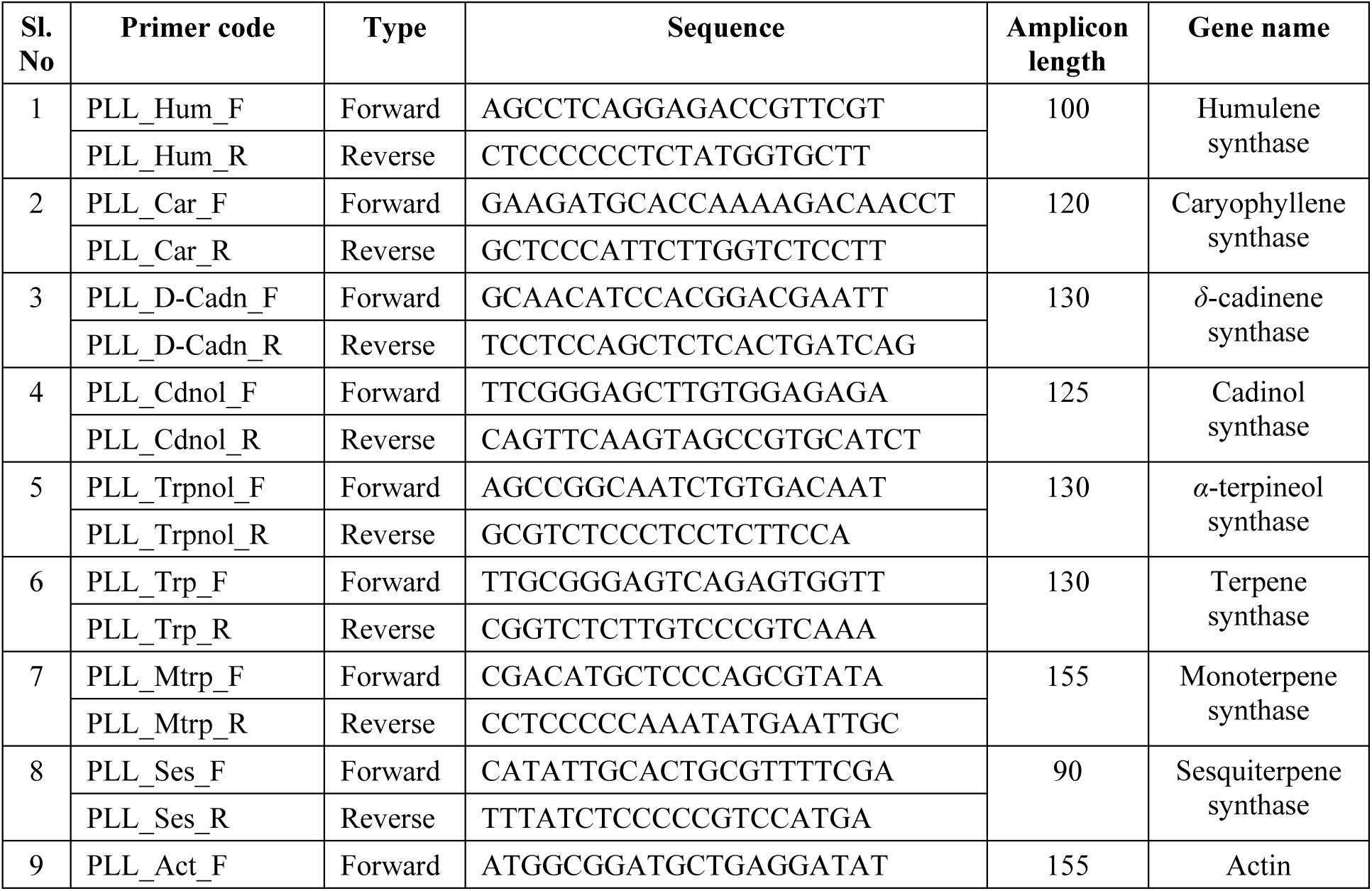
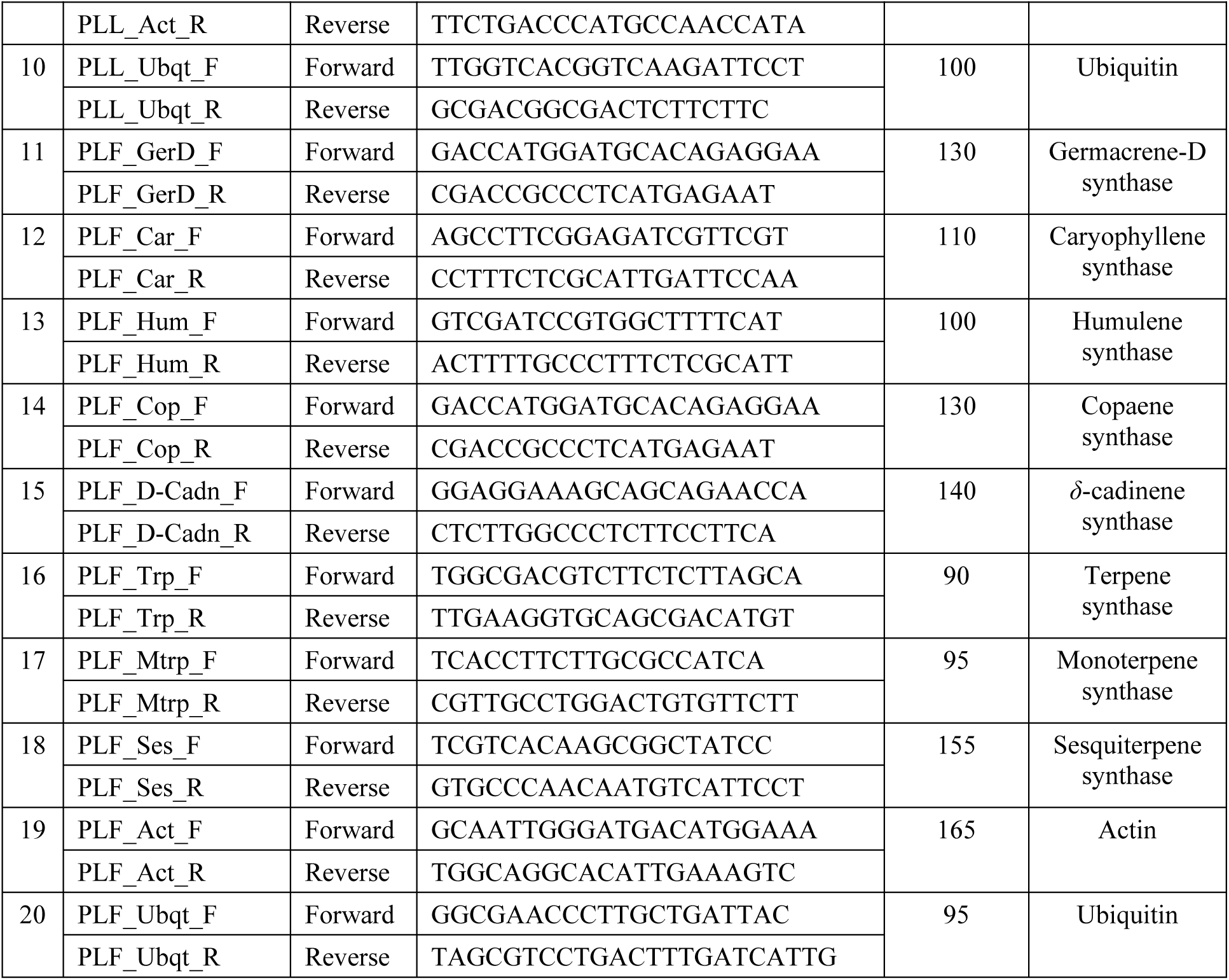
List of primers related to essential oil biosynthesis for RT-PCR analysis.

**Table 2:** Alignment statistics of *P*. *longum* RNA sequencing.

| Sl. No. | Sample | Total Reads | Processed Reads | %High quality |
| --- | --- | --- | --- | --- |
| 1 | Leaf | 36914662 | 36688460 | 99.39 |
| 2 | Fruit | 36244013 | 36011305 | 99.36 |

To perform *de novo* assembly, the clean reads were then assembled by RnaSPAdes software with a k-mer size of 55 by producing 1,90,596 (1.90 lakhs) and 2,03,110 (2.03 lakhs) reads with 972 and 891 bp N50 values for *P. longum* leaf and fruit respectively (Table 3). Maximum number of transcripts were of >=100-200 bp size with 55,231 and 54,852 transcripts followed by 54,349 and 52,819 transcripts of >=200-300 bp for leaf and fruit tissues respectively. In both tissues, least number of transcripts, which were 19 in leaf and 15 in fruit and were of >=10 kilo base pair (Kbp) to 1 mega base pair (Mbp). The average transcript lengths obtained to be 519.27.03 bp (leaf) and 517.04 bp (fruit) (Table 3).

**Table 3.** *De novo* assembly statistics of *P*. *longum* leaf and fruit.

|  | Leaf |  |  | Fruit |  |  |
| --- | --- | --- | --- | --- | --- | --- |
|  | Assembled | Clustered | CDS | Assembled | Clustered | CDS |
| Number of Sequences | 190596 | 78465 | 40033 | 203110 | 90836 | 44025 |
| Total Sequences Length | 98971083 | 77054596 | 29553447 | 105015114 | 82928025 | 31462371 |
| Maximum Sequence Length | 24985 | 24985 | 15276 | 16934 | 16934 | 15273 |
| Minimum Sequence Length | 43 | 300 | 297 | 40 | 300 | 297 |
| Average Sequence Length | 519.27 | 982.03 | 738.23 | 517.04 | 912.94 | 714.65 |
| Number of non-ATGC Characters | 42221 | 42211 | 66769 | 42970 | 42970 | 58555 |
| % of non-ATGC Characters | 0.0427 | 0.0548 | 0.2259 | 0.0409 | 0.0518 | 0.1861 |
| Sequences <100 bp | 1554 | 0 | 0 | 2740 | 0 | 0 |
| Sequences $\geq 100$-200 bp | 55231 | 0 | 0 | 54852 | 0 | 0 |
| Sequences $\geq 200$-300 bp | 54349 | 0 | 89 | 52819 | 0 | 117 |
| Sequences $\geq 300$-400 bp | 18338 | 17794 | 11317 | 22845 | 21673 | 12880 |
| Sequences $\geq 400$-500 bp | 11038 | 10765 | 6794 | 13845 | 13408 | 7798 |
| Sequences $\geq 500$-600 bp | 7804 | 7718 | 4486 | 9354 | 9226 | 4953 |
| Sequences $\geq 600$-700 bp | 5841 | 5809 | 3315 | 6992 | 6938 | 3497 |
| Sequences $\geq 700$-800 bp | 4642 | 4620 | 2325 | 5420 | 5400 | 2619 |
| Sequences $\geq 800$-900 bp | 3627 | 3614 | 1833 | 4420 | 4408 | 2008 |
| Sequences $\geq 900$-1Kbp | 3121 | 3114 | 1741 | 3651 | 3645 | 1879 |
| Sequences $\geq 1$-5Kbp | 24669 | 24649 | 8087 | 25843 | 25809 | 8240 |
| Sequences $\geq 5$-10Kbp | 363 | 363 | 40 | 314 | 314 | 29 |
| Sequences $\geq 10$Kb-1Mbp | 19 | 19 | 6 | 15 | 15 | 5 |
| N50 Value | 972 | 1391 | 894 | 891 | 1239 | 846 |

To eliminate the redundancy, library clusters and unigenes were obtained using the CD-HIT software which resulted in a total of 78,465 (leaf) and 90,836 (fruit) contigs with a total of 77,054,596 bp in leaf and 82,928,025 bp in fruit. The minimum sequence length was 300 bp both in leaf as well as fruit and maximum sequence length was 24,985 bp in leaf and 16,934 bp in fruit. In both the tissues, the average contig lengths were 982.03 bp and 912.94 bp with N50 values of 1391 and 1239 and bp in leaf and fruit respectively. The highest number of contigs were of >=1-5 Kbp with 25,809 transcripts in fruit and 24,649 in leaf (Table 3).

The TransDecoder tool was used to identify candidate coding regions in each transcript after clustering and rectifying the errors. A total of 40,033 and 44,025 CDS were predicted from the unigenes using having a total of 29,553,447 and 31,462,371 bases in leaf and fruit respectively. The mean CDS length in leaf and fruit were obtained to be 738.23 and 714.65 bp with a maximum length of 15,276 and 15,273 bp respectively (Figs. 2A and 2B).

**Fig 2.**
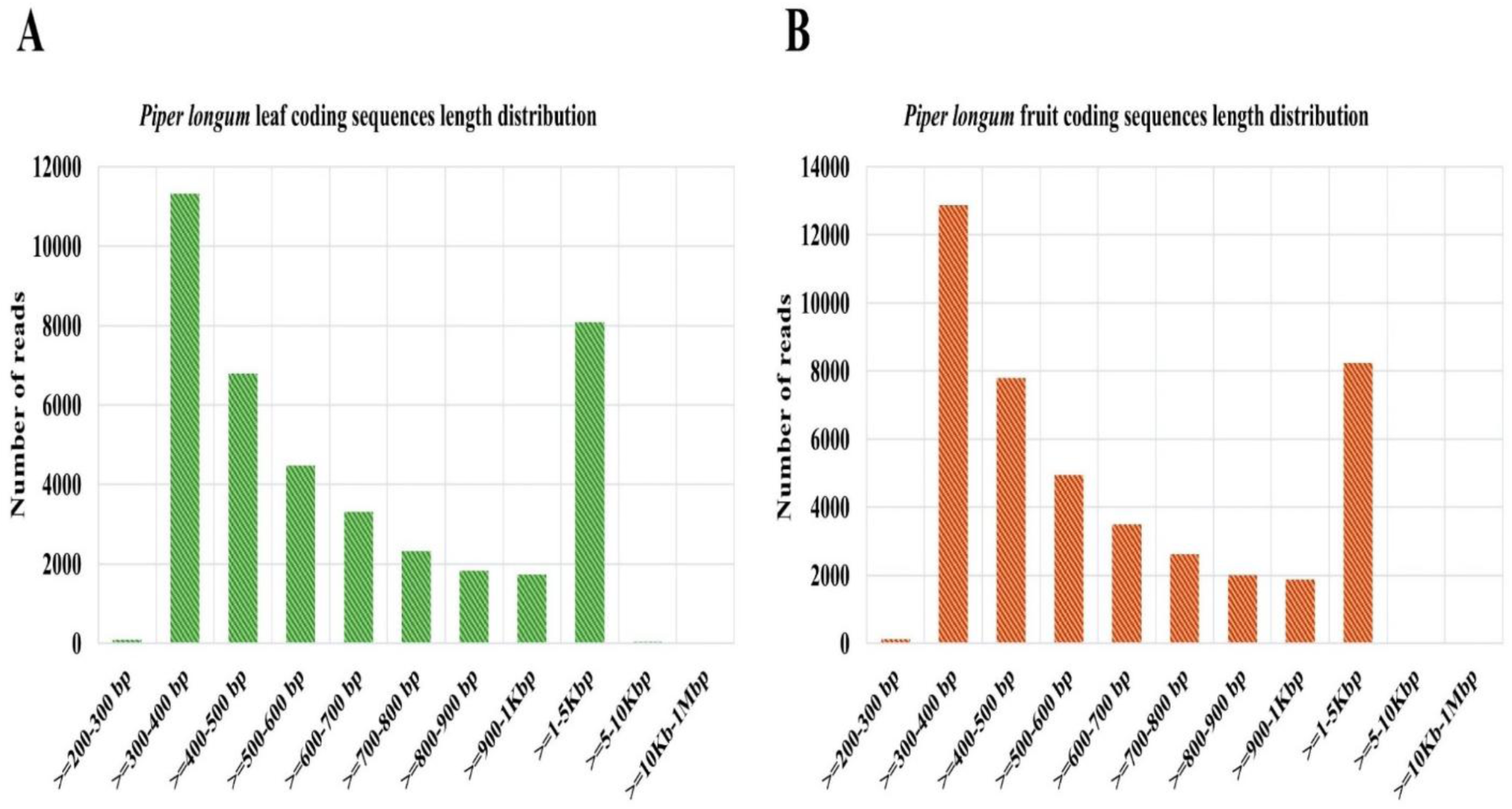
Length distribution of coding sequences of (A) leaf and (B) fruit of *P*. *longum*. Functional annotation of long pepper transcripts.

The Gene Ontology (GO) annotation was done using Diamond BLAST tool with default parameters by considering 40,033 and 44,025 transcripts of leaf and fruit respectively. Out of 40,033 and 44,025 transcripts, 34,045 (85.04%) and 37,037 (84.12%) transcripts have been annotated successfully in three different classes. GO analysis revealed that, class of cellular components was most abundant in both leaf (42.53%) and fruit (42.10%) samples followed by molecular function in leaf (41.45%) and fruit (41.46%). The class having lowest abundance is biological process in both leaf (16%) and fruit (16.47%) respectively.

The cellular components category mainly covered the cells, cell parts, organelles, membrane parts etc. whereas the molecular function category classified the gene into catalytic, binding activity, transporter activity etc. Lastly, the biological process consisted of metabolic process, cellular process, biological regulation etc. A detailed representation of GO annotation of both leaf and fruit were provided in Figs. 3A and 3B.

**Fig 3.**
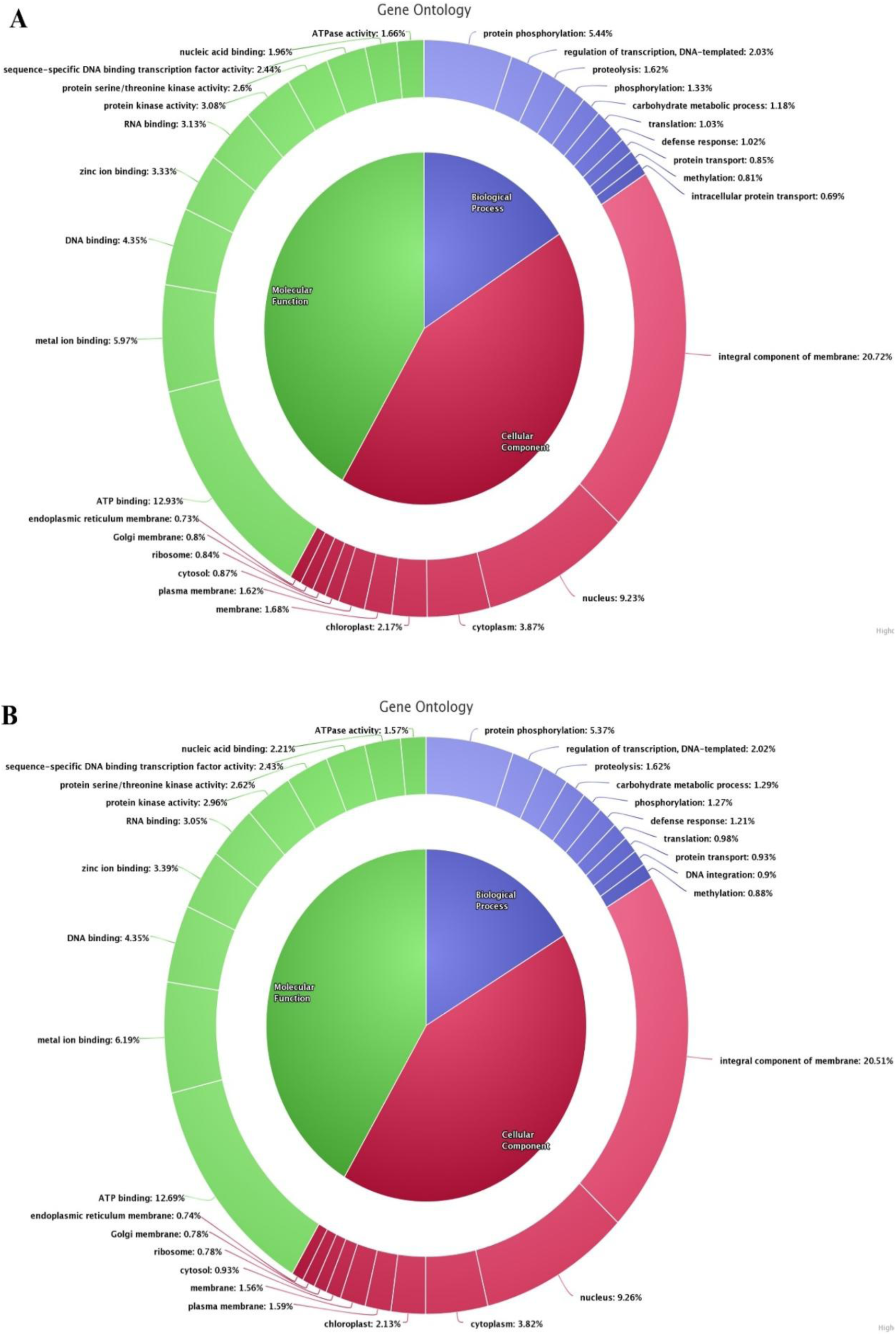
GO annotation showing cellular components, molecular function, and biological processes of (A) leaf and (B) fruit of *P*. *longum*.

### Identification of key genes involved in essential oil biosynthesis

From the GO annotation dataset, the genes involved in essential oil biosynthesis were also identified. The study mined 94 and 106 genes involved in terpenoid backbone biosynthesis comprises of both MVA and MEP/DXP pathways in leaf and fruit samples respectively. In *P. longum* leaf samples, 1-deoxy-D-xylulose-5-phosphate synthase (12) was found to be the frequently identified one followed by terpene synthase (10), (+)-δ-cadinene synthase (10), 4-hydroxy-3-methylbut-2-en-1-yl-diphosphate synthase (7), farnesyl pyrophosphate synthase (5), 3-hydroxy-3-methylglutaryl coenzyme A reductase (4), hydroxy methylglutaryl coenzyme A synthase (4), 4-hydroxy-3-methylbut-2-en-1-yl diphosphate reductase (4) sesquiterpene synthase (4), cadinol synthase (4), β-caryophyllene synthase (4), α-humulene synthase (4), (-)-α-terpineol synthase (3), diphosphomevalonate kinase (3), Isopentenyl-diphosphate δ-isomerase (3) and others which are figured out in Fig. 4A.

**Fig 4.**
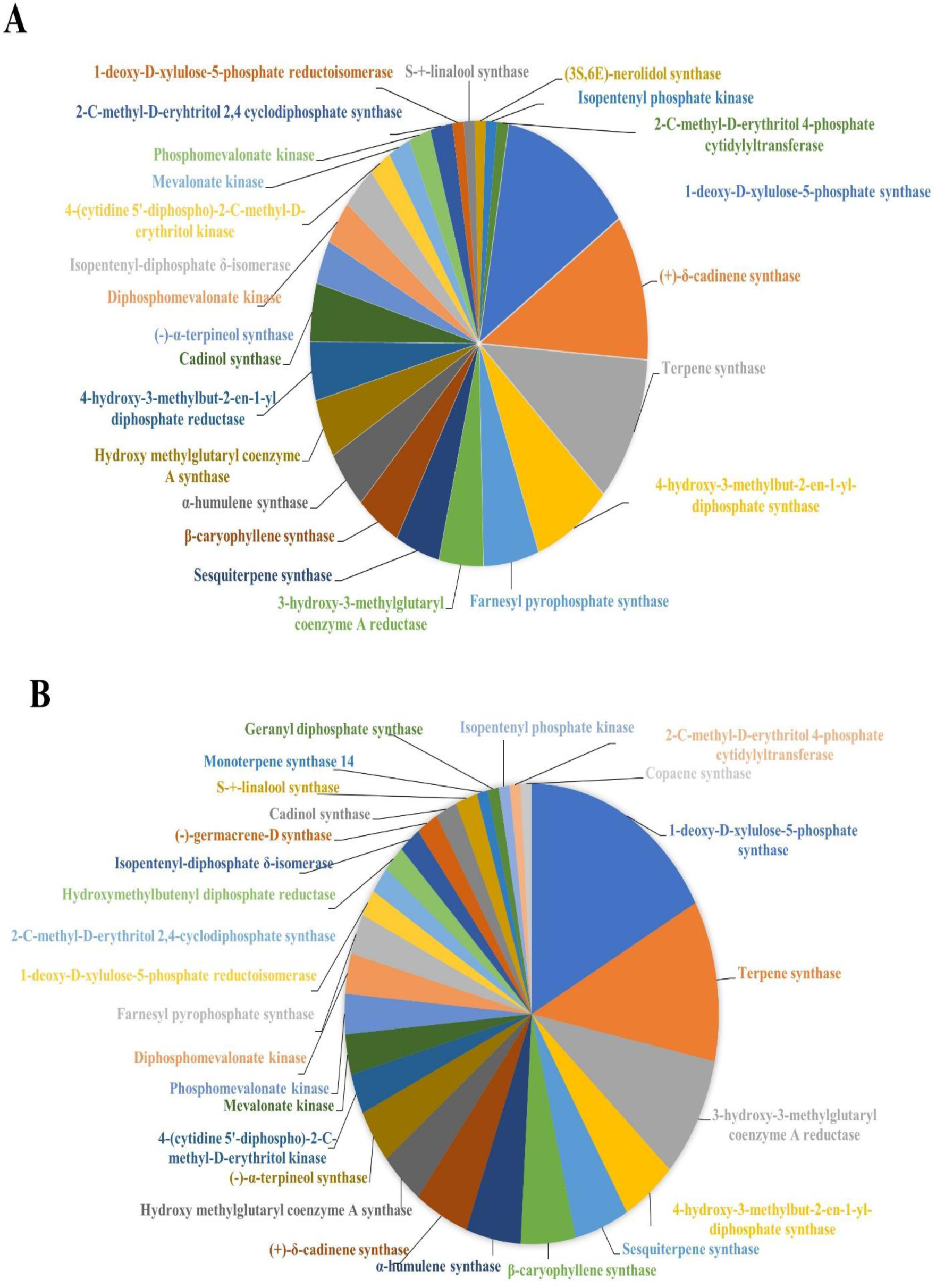
Mining of terpene biosynthesis genes from Gene Ontology; Identified terpene genes in (A) leaf, and (B) fruit of *P*. *longum*.

Similarly, in long pepper fruit sample, the most frequently detected gene was 1-deoxy-D-xylulose-5-phosphate synthase (18) followed by terpene synthase (12), 3-hydroxy-3-methylglutaryl coenzyme A reductase (9), 4-hydroxy-3-methylbut-2-en-1-yl-diphosphate synthase (5), sesquiterpene synthase (5), β-caryophyllene synthase (5), α-humulene synthase (5), (+)-δ-cadinene synthase (5), hydroxy methylglutaryl coenzyme A synthase (4), (-)-α-terpineol synthase (4), 4-(cytidine 5’-diphospho)-2-C-methyl-D-erythritol kinase (3), mevalonate kinase (3), phosphomevalonate kinase (3), diphosphomevalonate kinase (3), farnesyl pyrophosphate synthase (3), 1-deoxy-D-xylulose-5-phosphate reductoisomerase (2), 2-C-methyl-p-eryhtritol 2,4 cyclodiphosphate synthase (2), hydroxymethylbutenyl diphosphate reductase (2), isopentenyl-diphosphate δ-isomerase (2), (-)-germacrene-D synthase (2), cadinol synthase (2), S-+-linalool synthase (2) and some more were delineated in Fig. 4B.

### Annotation of transcripts using KEGG database

Annotation of pathways reveals the presence of multiple important secondary metabolite pathways that produce compounds with a wide range of therapeutic effects. Biochemical pathways associated with unigenes are determined using the KEGG database. The KEGG annotation resulted 29,166 candidate transcripts were involved in 203 different pathways identified in fruit sample whereas in leaf sample, 31,380 transcripts were assigned to 204 pathways. In leaf sample, a total count of 138 candidate transcripts (0.47%) belonging to three main categories were revealed having role in terpene synthase such as; terpenoid backbone biosynthesis (101) (73%), monoterpenoid biosynthesis (20) (15%), and sesquiterpenoid biosynthesis (17) (12%) (Fig. 5A). Likewise, in fruit sample, a total count of 155 transcripts (0.49%) were having potential role in terpene synthase distributed in three major categories, such as; terpenoid backbone biosynthesis (110) (71%), monoterpenoid biosynthesis (23) (15%), sesquiterpenoid biosynthesis (22) (14%) (Fig. 5B).

**Fig 5.**
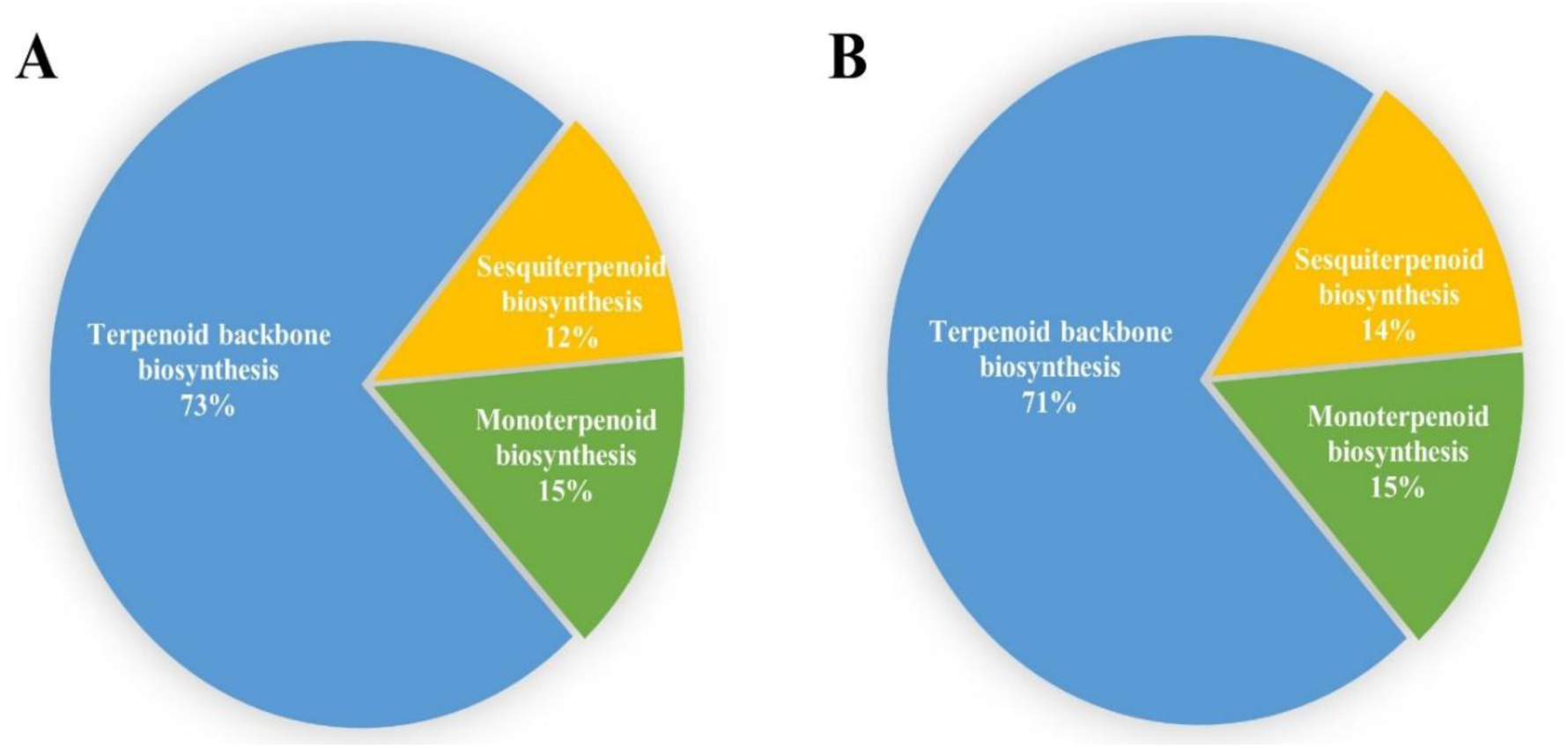
KEGG annotation showing the transcripts involved in terpinene biosynthesis of *P*. *longum* (A) leaf and (B) fruit.

### GC-MS analysis

The leaf and fruit EOs gave oil yield of and 0.15±0.028% and 0.88±0.04% based on the dry weight (v/w) of samples respectively. The obtained oils were then analysed through GC-MS towards the identification of terpenic composition. The constituents were identified by annotating the spectra with Adams and NIST library. The analysis revealed 53 and 58 constituents in leaf and fruit volatiles representing 97.99% and 98.91%, respectively. The identified constituents of leaf volatiles were categorized into monoterpene hydrocarbons, oxygenated monoterpenes, aldehyde, alkanes, ester, ether, ketones, sesquiterpene hydrocarbons, oxygenated sesquiterpenes. Similarly, the phytoconstituents identified in fruit volatiles were classified into monoterpene hydrocarbons, oxygenated monoterpenes, alcohols, alkanes, alkenes, esters, phenylpropanoid, sesquiterpene hydrocarbons and oxygenated sesquiterpenes. In leaf EO, monoterpene hydrocarbons were the predominant class in the leaf EOs with 34.46% followed by oxygenated sesquiterpenes (28.6%) and sesquiterpene hydrocarbons (18.27%). However, the fruit volatile composition predominated with sesquiterpene hydrocarbons with 53.53% followed by alkanes (14.81%) and alkenes (13.21%). Further, the analysis revealed *E*-nerolidol as the major constituents in the leaf EO with 21.59% followed by *β*-pinene (19.63%), and *α*-pinene (7.96%). However, the fruit EO dominated with germacrene-D (22.3%) followed by 8-heptadecene (10%), and *β*-caryophyllene (9.24%). The identified constituents of both leaf and fruit were summarised in Table 4 and the ion chromatograms marked with top 5 constituents were represented in Figs. 6A and 6B.

**Fig 6.**
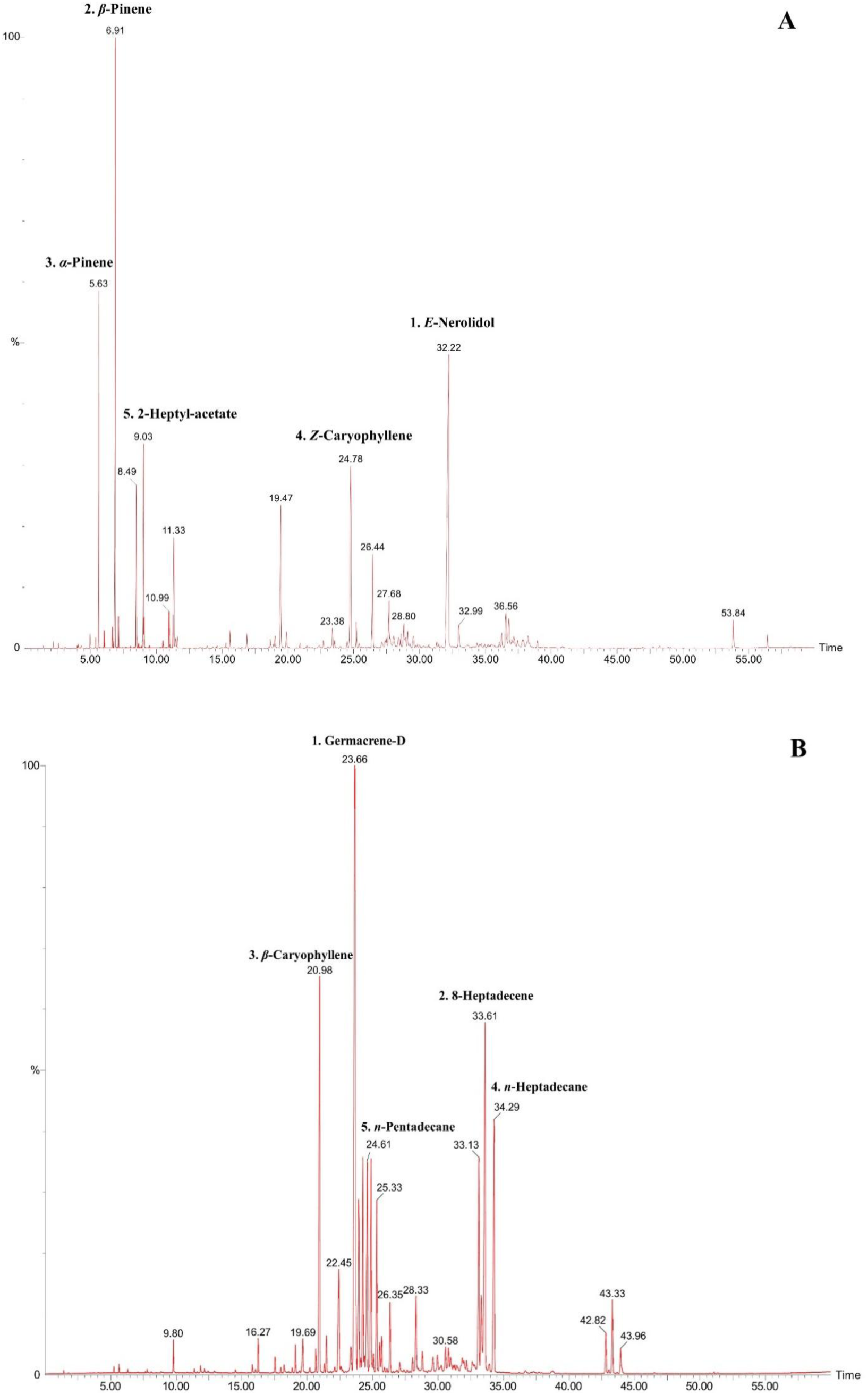
Chromatogram obtained from gas chromatography- mass spectrometry (GC-MS) analysis of (A) leaf and (B) fruit of *P*. *longum*.

**Table 4:**
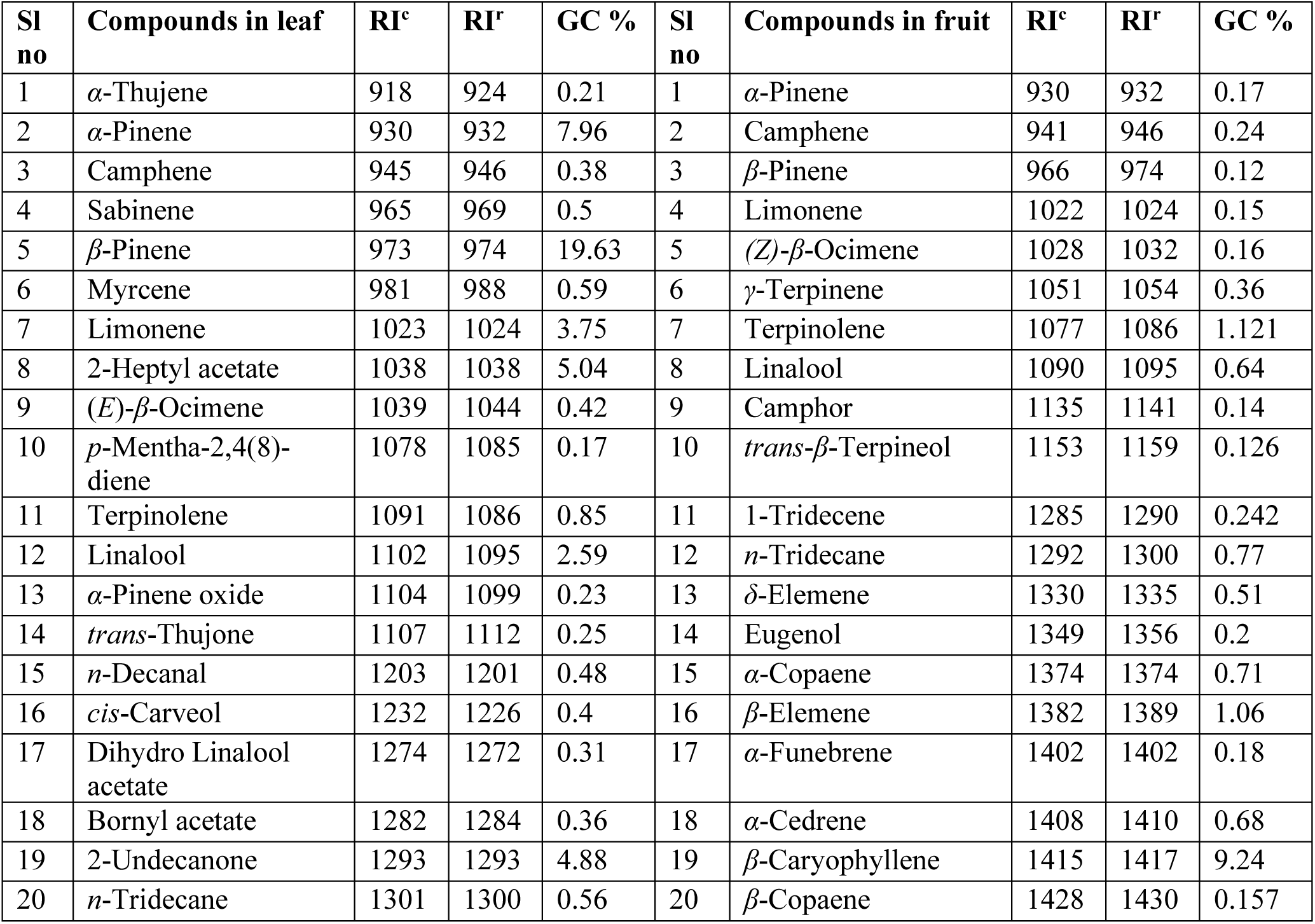
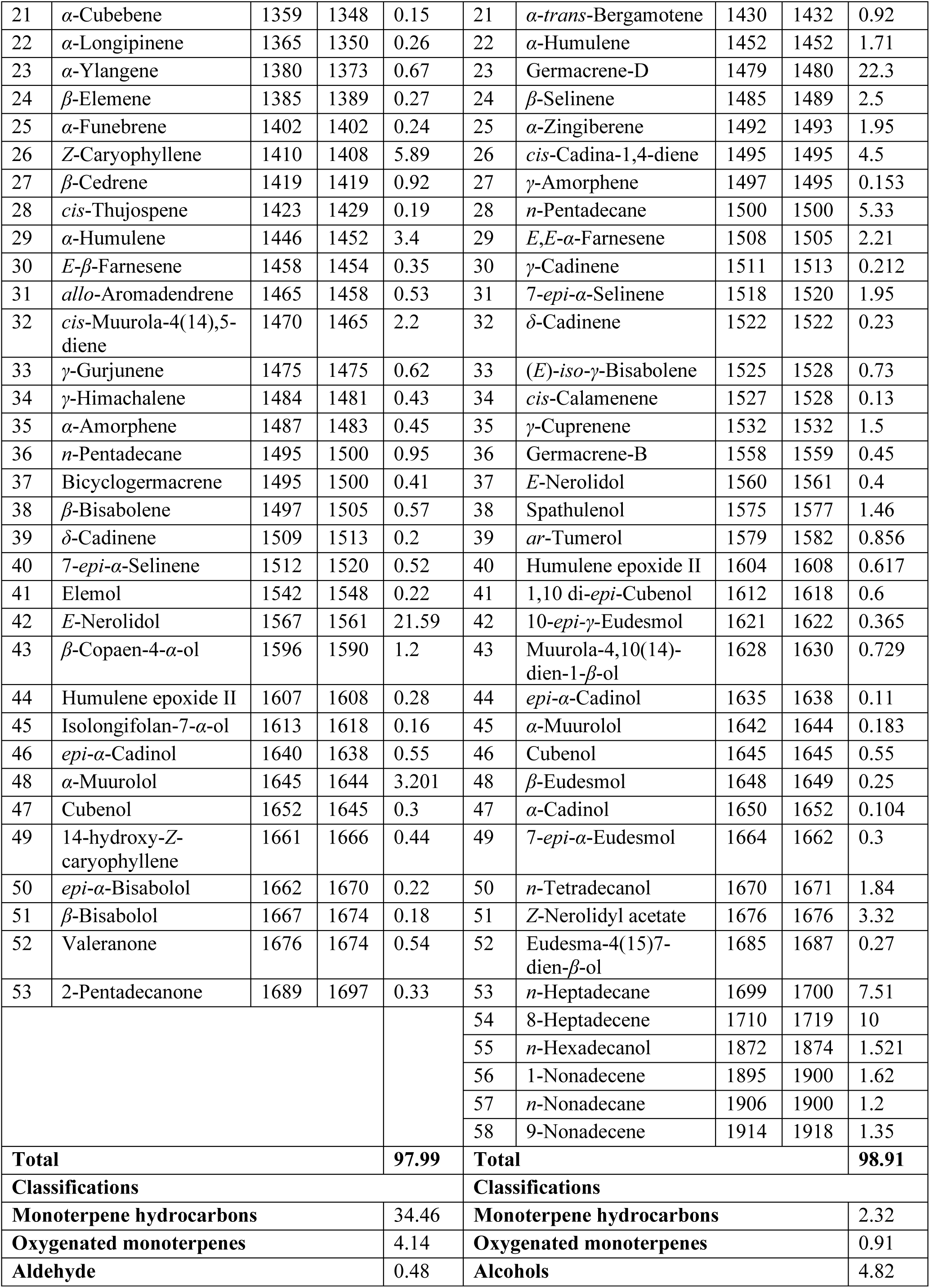
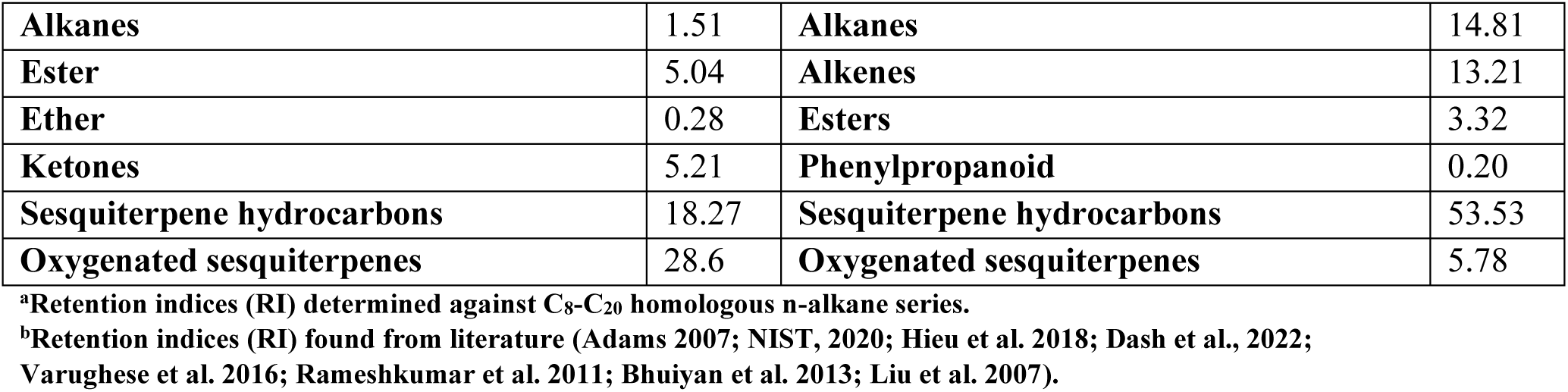
List of constituents identified by GC-MS in leaf and fruit of *P. longum*.

| Sl no | Compounds in leaf | RI<sup>c</sup> | RI<sup>r</sup> | GC % | Sl no | Compounds in fruit | RI<sup>c</sup> | RI<sup>r</sup> | GC % |
| --- | --- | --- | --- | --- | --- | --- | --- | --- | --- |
| 1 | α-Thujene | 918 | 924 | 0.21 | 1 | α-Pinene | 930 | 932 | 0.17 |
| 2 | α-Pinene | 930 | 932 | 7.96 | 2 | Camphene | 941 | 946 | 0.24 |
| 3 | Camphene | 945 | 946 | 0.38 | 3 | β-Pinene | 966 | 974 | 0.12 |
| 4 | Sabinene | 965 | 969 | 0.5 | 4 | Limonene | 1022 | 1024 | 0.15 |
| 5 | β-Pinene | 973 | 974 | 19.63 | 5 | (*Z*)-β-Ocimene | 1028 | 1032 | 0.16 |
| 6 | Myrcene | 981 | 988 | 0.59 | 6 | γ-Terpinene | 1051 | 1054 | 0.36 |
| 7 | Limonene | 1023 | 1024 | 3.75 | 7 | Terpinolene | 1077 | 1086 | 1.121 |
| 8 | 2-Heptyl acetate | 1038 | 1038 | 5.04 | 8 | Linalool | 1090 | 1095 | 0.64 |
| 9 | (*E*)-β-Ocimene | 1039 | 1044 | 0.42 | 9 | Camphor | 1135 | 1141 | 0.14 |
| 10 | *p*-Mentha-2,4(8)-diene | 1078 | 1085 | 0.17 | 10 | *trans*-β-Terpineol | 1153 | 1159 | 0.126 |
| 11 | Terpinolene | 1091 | 1086 | 0.85 | 11 | 1-Tridecene | 1285 | 1290 | 0.242 |
| 12 | Linalool | 1102 | 1095 | 2.59 | 12 | *n*-Tridecane | 1292 | 1300 | 0.77 |
| 13 | α-Pinene oxide | 1104 | 1099 | 0.23 | 13 | δ-Elemene | 1330 | 1335 | 0.51 |
| 14 | *trans*-Thujone | 1107 | 1112 | 0.25 | 14 | Eugenol | 1349 | 1356 | 0.2 |
| 15 | *n*-Decanal | 1203 | 1201 | 0.48 | 15 | α-Copaene | 1374 | 1374 | 0.71 |
| 16 | *cis*-Carveol | 1232 | 1226 | 0.4 | 16 | β-Elemene | 1382 | 1389 | 1.06 |
| 17 | Dihydro Linalool acetate | 1274 | 1272 | 0.31 | 17 | α-Funebrene | 1402 | 1402 | 0.18 |
| 18 | Bornyl acetate | 1282 | 1284 | 0.36 | 18 | α-Cedrene | 1408 | 1410 | 0.68 |
| 19 | 2-Undecanone | 1293 | 1293 | 4.88 | 19 | β-Caryophyllene | 1415 | 1417 | 9.24 |
| 20 | *n*-Tridecane | 1301 | 1300 | 0.56 | 20 | β-Copaene | 1428 | 1430 | 0.157 |
| 21 | α-Cubebene | 1359 | 1348 | 0.15 | 21 | α-trans-Bergamotene | 1430 | 1432 | 0.92 |
| 22 | α-Longipinene | 1365 | 1350 | 0.26 | 22 | α-Humulene | 1452 | 1452 | 1.71 |
| 23 | α-Ylangene | 1380 | 1373 | 0.67 | 23 | Germacrene-D | 1479 | 1480 | 22.3 |
| 24 | β-Elemene | 1385 | 1389 | 0.27 | 24 | β-Selinene | 1485 | 1489 | 2.5 |
| 25 | α-Funebrene | 1402 | 1402 | 0.24 | 25 | α-Zingiberene | 1492 | 1493 | 1.95 |
| 26 | Z-Caryophyllene | 1410 | 1408 | 5.89 | 26 | cis-Cadina-1,4-diene | 1495 | 1495 | 4.5 |
| 27 | β-Cedrene | 1419 | 1419 | 0.92 | 27 | γ-Amorphene | 1497 | 1495 | 0.153 |
| 28 | cis-Thujospene | 1423 | 1429 | 0.19 | 28 | n-Pentadecane | 1500 | 1500 | 5.33 |
| 29 | α-Humulene | 1446 | 1452 | 3.4 | 29 | E,E-α-Farnesene | 1508 | 1505 | 2.21 |
| 30 | E-β-Farnesene | 1458 | 1454 | 0.35 | 30 | γ-Cadinene | 1511 | 1513 | 0.212 |
| 31 | allo-Aromadendrene | 1465 | 1458 | 0.53 | 31 | 7-epi-α-Selinene | 1518 | 1520 | 1.95 |
| 32 | cis-Muurolo-4(14),5-diene | 1470 | 1465 | 2.2 | 32 | δ-Cadinene | 1522 | 1522 | 0.23 |
| 33 | γ-Gurjunene | 1475 | 1475 | 0.62 | 33 | (E)-iso-γ-Bisabolene | 1525 | 1528 | 0.73 |
| 34 | γ-Himachalene | 1484 | 1481 | 0.43 | 34 | cis-Calamenene | 1527 | 1528 | 0.13 |
| 35 | α-Amorphene | 1487 | 1483 | 0.45 | 35 | γ-Cuprenene | 1532 | 1532 | 1.5 |
| 36 | n-Pentadecane | 1495 | 1500 | 0.95 | 36 | Germacrene-B | 1558 | 1559 | 0.45 |
| 37 | Bicyclogermacrene | 1495 | 1500 | 0.41 | 37 | E-Nerolidol | 1560 | 1561 | 0.4 |
| 38 | β-Bisabolene | 1497 | 1505 | 0.57 | 38 | Spathulenol | 1575 | 1577 | 1.46 |
| 39 | δ-Cadinene | 1509 | 1513 | 0.2 | 39 | ar-Tumerol | 1579 | 1582 | 0.856 |
| 40 | 7-epi-α-Selinene | 1512 | 1520 | 0.52 | 40 | Humulene epoxide II | 1604 | 1608 | 0.617 |
| 41 | Elemol | 1542 | 1548 | 0.22 | 41 | 1,10 di-epi-Cubenol | 1612 | 1618 | 0.6 |
| 42 | E-Nerolidol | 1567 | 1561 | 21.59 | 42 | 10-epi-γ-Eudesmol | 1621 | 1622 | 0.365 |
| 43 | β-Copaen-4-α-ol | 1596 | 1590 | 1.2 | 43 | Muurolo-4,10(14)-dien-1-β-ol | 1628 | 1630 | 0.729 |
| 44 | Humulene epoxide II | 1607 | 1608 | 0.28 | 44 | epi-α-Cadinol | 1635 | 1638 | 0.11 |
| 45 | Isolongifolan-7-α-ol | 1613 | 1618 | 0.16 | 45 | α-Muurolol | 1642 | 1644 | 0.183 |
| 46 | epi-α-Cadinol | 1640 | 1638 | 0.55 | 46 | Cubenol | 1645 | 1645 | 0.55 |
| 48 | α-Muurolol | 1645 | 1644 | 3.201 | 48 | β-Eudesmol | 1648 | 1649 | 0.25 |
| 47 | Cubenol | 1652 | 1645 | 0.3 | 47 | α-Cadinol | 1650 | 1652 | 0.104 |
| 49 | 14-hydroxy-Z-caryophyllene | 1661 | 1666 | 0.44 | 49 | 7-epi-α-Eudesmol | 1664 | 1662 | 0.3 |
| 50 | epi-α-Bisabolol | 1662 | 1670 | 0.22 | 50 | n-Tetradecanol | 1670 | 1671 | 1.84 |
| 51 | β-Bisabolol | 1667 | 1674 | 0.18 | 51 | Z-Nerolidyl acetate | 1676 | 1676 | 3.32 |
| 52 | Valeranone | 1676 | 1674 | 0.54 | 52 | Eudesma-4(15)7-dien-β-ol | 1685 | 1687 | 0.27 |
| 53 | 2-Pentadecanone | 1689 | 1697 | 0.33 | 53 | n-Heptadecane | 1699 | 1700 | 7.51 |
|  |  |  |  |  | 54 | 8-Heptadecene | 1710 | 1719 | 10 |
|  |  |  |  |  | 55 | n-Hexadecanol | 1872 | 1874 | 1.521 |
|  |  |  |  |  | 56 | 1-Nonadecene | 1895 | 1900 | 1.62 |
|  |  |  |  |  | 57 | n-Nonadecane | 1906 | 1900 | 1.2 |
|  |  |  |  |  | 58 | 9-Nonadecene | 1914 | 1918 | 1.35 |
| **Total** |  |  |  | **97.99** | **Total** |  |  |  | **98.91** |
| **Classifications** |  |  |  |  | **Classifications** |  |  |  |  |
| **Monoterpene hydrocarbons** |  |  |  | 34.46 | **Monoterpene hydrocarbons** |  |  |  | 2.32 |
| **Oxygenated monoterpenes** |  |  |  | 4.14 | **Oxygenated monoterpenes** |  |  |  | 0.91 |
| **Aldehyde** |  |  |  | 0.48 | **Alcohols** |  |  |  | 4.82 |

|  |  |  |  |
| --- | --- | --- | --- |
| **Alkanes** | 1.51 | **Alkanes** | 14.81 |
| **Ester** | 5.04 | **Alkenes** | 13.21 |
| **Ether** | 0.28 | **Esters** | 3.32 |
| **Ketones** | 5.21 | **Phenylpropanoid** | 0.20 |
| **Sesquiterpene hydrocarbons** | 18.27 | **Sesquiterpene hydrocarbons** | 53.53 |
| **Oxygenated sesquiterpenes** | 28.6 | **Oxygenated sesquiterpenes** | 5.78 |

<sup>a</sup>Retention indices (RI) determined against C<sub>8</sub>-C<sub>20</sub> homologous n-alkane series.
<sup>b</sup>Retention indices (RI) found from literature (Adams 2007; NIST, 2020; Hieu et al. 2018; Dash et al., 2022; Varughese et al. 2016; Rameshkumar et al. 2011; Bhuiyan et al. 2013; Liu et al. 2007).

### Identification of SSRs

MISA tool examined 38,467 and 42,268 number of sequences with total size of 28574751 and 30428283 bp in long pepper leaf and fruit respectively. The total number of SSRs identified in leaf and fruit samples were 4086 and 4564 respectively. In leaf sample, 489 number of sequences identified containing more than 1 SSR and number of compound SSRs were 352. The fruit of *P. longum* found to have 539 number of sequences containing more than 1 SSR and 376 number of compound SSRs. The trinucleotide repeats were observed to be the most frequent occurring SSR motifs in leaf (2161) and fruit (2352) samples respectively followed by mononucleotide, tetranucleotide, dinucleotide, hexanucleotide, and pentanucleotide motifs in both the samples (Table 5, Figs. 7A and 7B). Among the identified SSR motifs, the T/A mononucleotide repeat motif was the most abundant in both leaf (323) and fruit (403) and samples followed by 205 numbers of CCG/GGC trinucleotide repeats in leaf samples and 209 in fruit samples respectively. The detailed information on SSR repeat motifs was represented in Figs. 7C and 7D.

**Fig 7.**
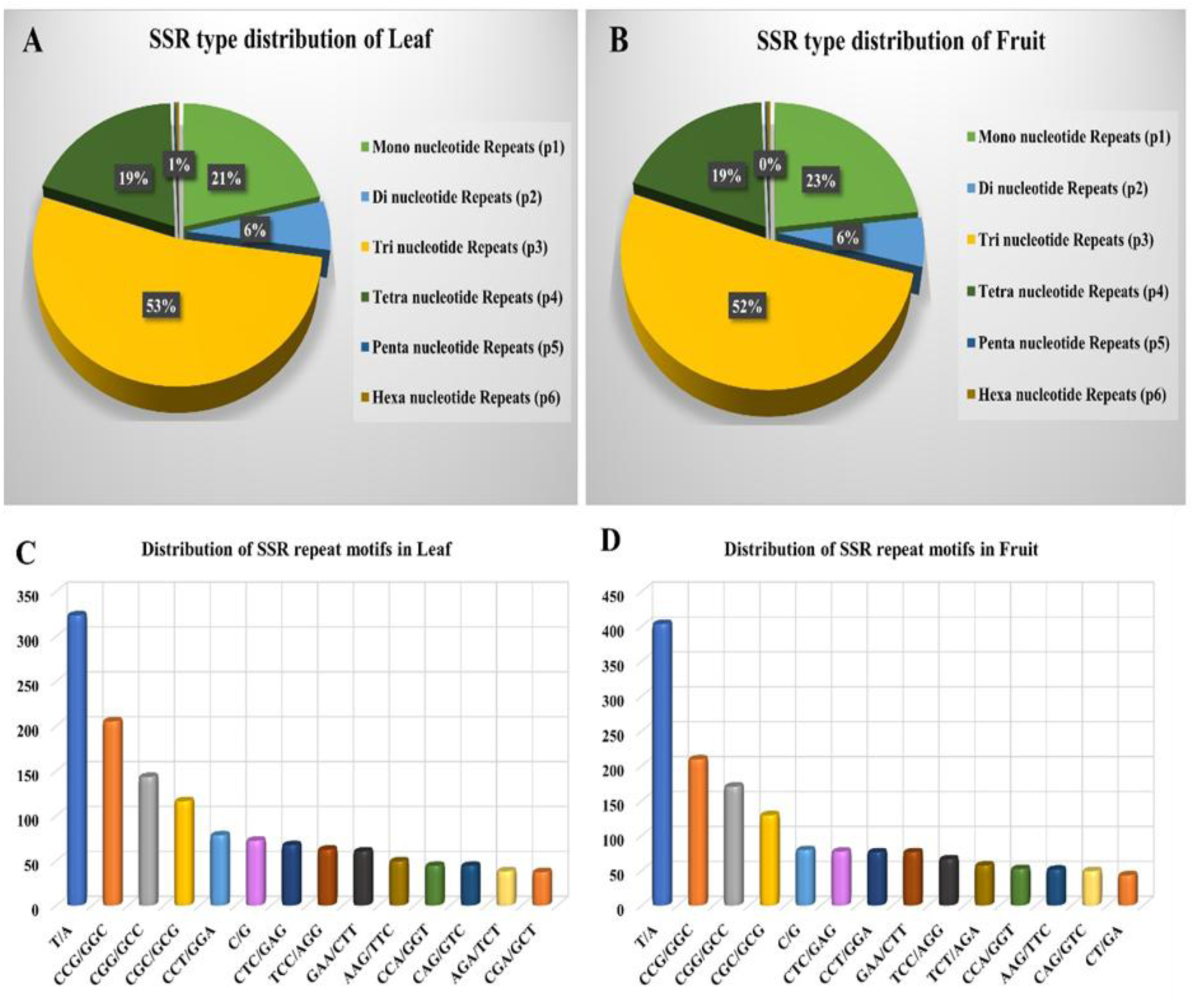
Distribution of SSR repeat type (A) leaf and (B) fruit; Distribution of SSR repeat motif (C) leaf and (D) fruit of *P*. *longum*.

**Table 5:**
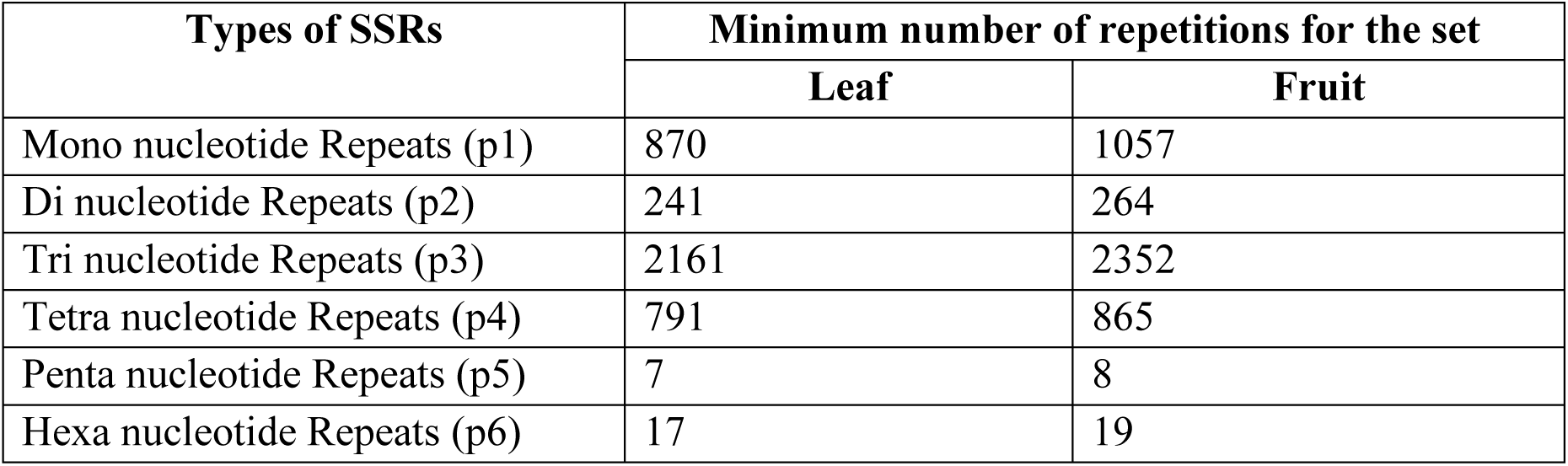
SSR statistics identified from transcripts of leaf and fruit of *P*. *longum*.

| Types of SSRs | Minimum number of repetitions for the set |  |
| --- | --- | --- |
|  | Leaf | Fruit |
| Mono nucleotide Repeats (p1) | 870 | 1057 |
| Di nucleotide Repeats (p2) | 241 | 264 |
| Tri nucleotide Repeats (p3) | 2161 | 2352 |
| Tetra nucleotide Repeats (p4) | 791 | 865 |
| Penta nucleotide Repeats (p5) | 7 | 8 |
| Hexa nucleotide Repeats (p6) | 17 | 19 |

### Relative gene expression study of terpenoid biosynthesis genes

The extracted good quality RNAs were subjected to cDNA synthesis. A total of eighteen number of transcript specific essential oil biosynthesis primers were studied for relative gene expression in both leaf and fruit samples. The experiment was conducted in triplicates (Figs. 8A and 8B). The terpene synthase gene in leaf sample has showed a higher level of expression followed by monoterpene and sesquiterpene synthases. Similarly, in fruit samples, terpene synthase gene showed higher expression followed by sesquiterpene synthase and monoterpene synthase genes. However, among the other transcript specific primers, the expression of caryophyllene synthase gene was higher followed by humulene synthase gene and *δ*-cadinene synthase genes in leaf. In fruit sample, germacrene-D followed by caryophyllene synthase genes has shown higher expression than other genes. In both the samples, humulene synthase, copaene synthase, *δ*-cadinene synthase and cadinol synthase genes also showed their expression (Figs. 8A and 8B). The *α*-terpineol synthase gene has shown the least expression leaf. These terpene biosynthesis genes were also present in varying area percentages in long pepper leaf and fruit samples and GC-MS analysis was implemented to further identify and analyze them.

**Fig 8.**
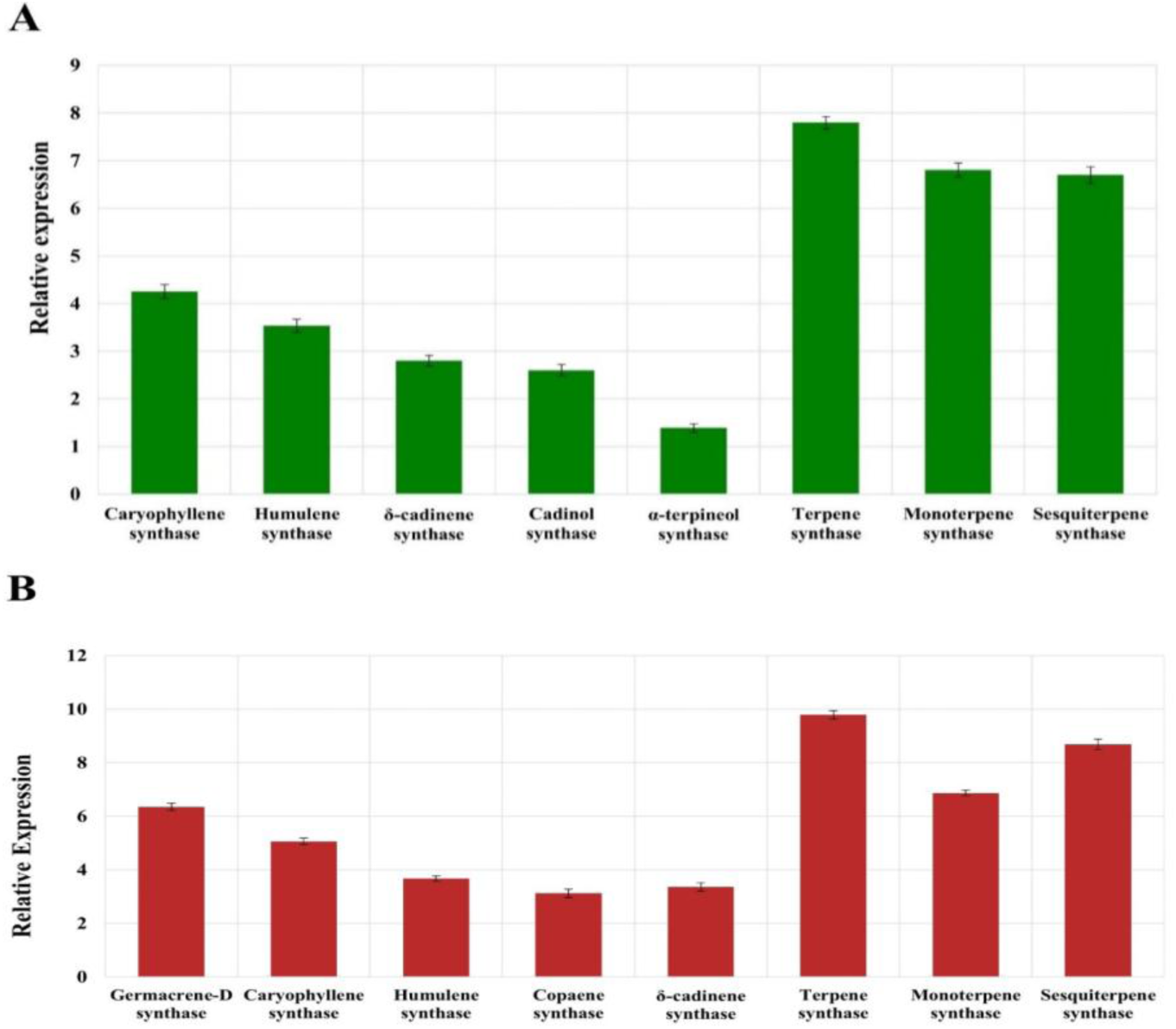
Terpene biosynthesis gene expression using RT-PCR in long pepper (A) leaf and (B) fruit.

## Discussion

Long pepper is one of the most significant medicinal plants possessing wide range of medicinal uses. This plant is frequently used in treating several diseases. India is one of the important hubs for cultivating long pepper and exporting it in substantial quantities to nations around the globe. This species not only has extensive medicinal importance, but also being utilized for generating livelihoods. The pharmacological properties of long pepper are mainly reliant on secondary metabolites. As previously stated, the terpenoid contents of this species exhibits a variety of medicinal properties. Thus, understanding the molecular processes underlying these terpenoid synthesis is crucial for the taxa’s future development. One of the efficient strategies to identify the terpenoid biosynthesis pathway is transcriptome analysis, which has been studied in numerous non-model plants like *Piper betle* (Sahoo et al., 2021), *Piper nigrum* (Hao et al., 2016), *Litsea cubeba* (Han et al., 2013), *Gymnema sylvestre* (Ayachit et al., 2019), *Salvia hispanica* (Wimberley et al., 2020), *Salvia guaraniti*ca (Ali et al., 2018), *Cinnamomum camphora* (Chen et al., 2018), *Curcuma longa* (Sahoo et al., 2019), *Ocimum sanctum* and *Ocimum basilicum* (Rastogi et al., 2014).

In the present study, illumina platform was employed for whole RNA sequencing towards studying the terpenoid biosynthesis pathway.

In the present study, illumina platform was employed for whole RNA sequencing towards studying the terpenoid biosynthesis pathway. Prior to assembly, low quality reads and adapters must be removed from the raw data. The N50 value is another key factor responsible for establishing quality assembly. In this study, the *de novo* transcriptome of long pepper generated transcripts having average contig length of 912.94 and 982.03 bp with N50 values of 1239 and 1391 bp in leaf and fruit samples, respectively. The obtained result was almost similar to the previously reported *de novo* transcriptome analysis of *P. nigrum* (Hu et al., 2015), *G. sylvestre* (Ayachit et al., 2019), *C. longa* (Sahoo et al., 2019), *L. cubeba* (Han et al., 2013), *O. sanctum* and *O. basilicum* (Rastogi et al., 2014). The N50 values were significantly better, so the generated high quality transcripts can be used for more in-depth analysis.

The transcripts were annotated using few databases like Diamond BLAST, GO, KEGG in order to analyze virtual functional characterization. The GO annotation of long pepper leaf and fruit were distributed into three classes; 42.53% and 42.10% transcripts were the most abundant in cellular components in leaf and fruit samples, respectively, 41.45% (leaf) and 41.46% (fruit) transcripts in molecular function. The class having lowest abundance is biological process in both leaf (16%) and fruit (16.47%) respectively. This result was nearly similar to the ontology annotation of *S. hispanica* (Wimberley et al., 2020) and *C. pictus* (Annadurai et al., 2012).

The GO annotation mined 94 and 106 genes involved in terpenoid backbone biosynthesis comprises of both MVA and MEP/DXP pathways in leaf and fruit samples respectively. Further, other terpenoid synthase genes were also identified which regulates monoterpenes, sesquiterpenes and diterpenes through both MVA and MEP/DXP pathways (Rastogi et al., 2014). In KEGG annotation, a total count of 138 candidate transcripts (0.47%) in leaf samples and 155 transcripts (0.49%) in fruit samples were having potential role in EO biosynthesis pathways. In both leaf and fruit samples, the maximum transcripts were found to be involved in terpenoid backbone biosynthesis. This result may be related to the phytochemical analysis of *P. longum* leaf and fruit EOs performed earlier using GC-MS, where majorly monoterpene and sesquiterpene classes of constituents were identified. The functional characterization of *P. longum* revealed that the RNA-Seq based *de novo* transcriptome analysis will further promote studies on the physiology, biochemistry, and molecular genetics of the species.

The relative gene expression study was conducted using seventeen transcript-specific primers both with leaf and fruit samples. In both the samples, terpene synthase gene was found with higher expression. In leaf samples, the next expressed gene is monoterpene synthase followed by sesquiterpene synthase genes. However, in fruit, the expression of sesquiterpene synthase gene was higher than monoterpene synthase gene. In leaf samples, caryophyllene synthase gene was found with relatively high expression followed by humulene synthase gene. However, in fruit germacrene-D synthase gene showed good expression followed by caryophyllene synthase gene. The obtained result was confirmed by chromatographic analyses of EOs. The GC-MS analysis showed monoterpene hydrocarbons, oxygenated sesquiterpene and sesquiterpene hydrocarbon classes of compounds possessed noticeable area percentage in leaf as well as fruit volatiles. It also revealed that the sesquiterpene hydrocarbons dominated the fruit volatiles with major constituent germacrene-D, and the third constituent is *β*-caryophyllene. The experimentally validated gene expression data will provide a better insight of function and regulation of genes.

In *P. longum*, there are no reports on the SSRs, which create difficulties in species identification. Generally, high quality polymorphic SSRs have putative role in comparative genomics, gene mapping, relatedness, linkage, breeding, population genetics, and genetic diversity and discrimination studies (Filippi et al., 2015, Mammadov et al., 2012, Qi and Ma, 2020, Singh et al., 2013, Hu et al., 2015). The current transcriptomic study revealed 4086 and 4564 numbers of SSRs in leaf and fruit samples, respectively using MISA tool. Trinucleotide repeats were found to be the dominant one both in leaf and fruit samples which corroborates with one of the previously reported studies by Hu et al., 2015 in fruits of *P. nigrum* (Hu et al., 2015) and *C. pictus* (Annadurai et al., 2012). In trinucleotide repeats, CGG/GCC in leaf and CCG/GGC in fruit were identified as the most abundant motifs. In this study, the genome wide SSRs identified in *P. longum* were discovered for the first time. This will strengthen the species’ molecular resources and help in future investigations towards genetic diversity, genetic linkage, and marker assisted selection for conventional plant breeding.

## Conclusions

Long pepper, a vital spice crop of India and has been used in Ayurvedic medicine for centuries due to its medicinal properties. To enhance its contribution to the national economy, it requires increased attention and investment. Despite its significance, long pepper has been limited by insufficient molecular resources for comprehensive genetic studies. Transcriptome sequencing is an efficient technique for identifying large-scale sequence information and has been employed in this study as a pioneering exploration of long pepper’s molecular resources. This study is the first attempt to explore molecular resources of *P. longum* through transcriptomics approach. he abundance of transcripts generated provides valuable insights into genes involved in various biochemical pathways, particularly terpenoid biosynthesis. Additionally, the dataset has facilitated the discovery of SSR markers. The present work may help functional genomics study of long pepper for its future improvement. In *P. longum*, the terpenoid biosynthesis pathway can be targeted for over production of desired industrially important compounds. This study will definitely attract long pepper growers, related industries and scientific communities as well, because of enormous information at phytochemical and molecular level.

## CRediT authorship contribution statement

Conceptualization: Basudeba Kar; Methodology: Basudeba Kar, Manaswini Dash, Bhaskar Chandra Sahoo; Material collection and processing: Manaswini Dash; Preparation of Figures: Manaswini Dash; Original draft preparation: Manaswini Dash, Bhaskar Chandra Sahoo; Manuscript writing: Manaswini Dash; Data interpretation: Manaswini Dash, Suprava Sahoo and Bhaskar Chandra Sahoo; Article review and editing: Basudeba Kar and Suprava Sahoo. All the authors have reviewed the manuscript.

## Acknowledgements

The authors thank Prof. (Dr.) Manoj Ranjan Nayak, Founder and President, Siksha O Anusandhan Deemed to be University for providing all the necessary facilities and encouragement as well. The authors also acknowledge the taxonomist Prof. P.C. Panda, Siksha O Anusandhan Deemed to be University for authenticating the sample.

## Data availability

The datasets generated and/or analyzed during the current study are available from the corresponding author on reasonable request.

## Competing interests

The authors declare that they do not have any conflict of interest.

## Notes

### Competing Interest Statement

The authors have declared no competing interest.

